# Anti-cancer compound screening identifies Aurora Kinase A inhibition as a means to favor CRISPR/Cas9 gene correction over knock-out

**DOI:** 10.1101/2023.11.09.566375

**Authors:** Danny Wilbie, Selma Eising, Vicky Amo-Addae, Johanna Walther, Esmeralda Bosman, Olivier G de Jong, Jan J Molenaar, Enrico Mastrobattista

## Abstract

CRISPR gene therapy holds the potential to cure a variety of genetic diseases by targeting causative mutations and introducing double stranded DNA breaks, subsequently allowing the host DNA repair mechanisms to introduce mutations. One option to introduce precise gene corrections is via the homology-directed repair (HDR) pathway. HDR can introduce a range of desired mutations dictated by a DNA template which holds a corrected DNA sequence which is written into the targeted gene. The problem in utilizing this pathway is that CRISPR-induced double stranded DNA breaks are repaired more often through the non-homologous end joining (NHEJ) pathway, which does not use a designed template and introduces random DNA damage in the form of insertions and deletions at the cut site. Since HDR activation depends on many interconnected processes in the cell, we aimed to screen a small library of drug compounds in clinical use or clinical development for cancer, to steer the DNA repair process towards preferential HDR activation.

We included compounds in our screen based on three relevant mechanisms in CRISPR gene editing: the cell cycle, DNA repair processing and chromosomal packing. We included forty compounds, based on these criteria, screened their toxicity and dosed them in sub-toxic concentrations in cells during genome editing. Of these forty compounds we identified nine potential hits to have an effect on preferential activation of the HDR pathway over NHEJ. Alisertib, rucaparib and belinostat revealed a significant and major effect on gene editing pathway selection in further validation.

Alisertib, an Aurora kinase A inhibitor, showed a particularly strong effect towards improving HDR over NHEJ. We subsequently investigated this effect at the genetic level and in a murine hepatoma cell line, which corroborated the initial findings. Alisertib led to an over 4-fold increase in preferential gene correction over gene knock-out, at a dose of 0.3 micromolar. However, the observations that Aurora kinase A inhibitors show considerable cytotoxicity (<50% cell viability) and can induce morphological changes at this concentration pose a limitation for the direct use of these inhibitors as HDR enhancers. However these findings do implicate that the pathways mediated by Aurora kinase A strongly influence HDR outcomes, which warrants further investigation into the downstream pathways driving this effect.

## Introduction

Curative gene editing by RNA-guided CRISPR/Cas9 nucleases has progressed into clinical trials in recent years, with applications varying from *ex vivo* cell modification to *in vivo* gene editing (1,2). This gene therapy method utilizes the CRISPR-associated protein 9 (Cas9) endonuclease, an enzyme which complexes with a guide RNA sequence which can direct it to a specific DNA target. This ribonucleoprotein complex binds to the DNA sequence complementary to the guide RNA, after which the Cas9 nuclease causes a double stranded DNA break (DSB). In the context of gene editing in cells, the RNP needs to reach the cell nucleus and bind to its target in the genomic DNA. Cells are equipped with DNA repair pathways to resolve this damage, which can be exploited for therapy (3).

Double stranded DNA breaks are predominately repaired by the non-homologous end-joining (NHEJ) and homology directed repair (HDR) pathways (4). NHEJ leads to ligation of the broken DNA ends by DNA ligase 4, which is often repaired perfectly but can also introduce small insertions or deletions (indels) at the double stranded break site. When repaired faithfully, this ligation leads to recovery of the Cas9 target sequence, which results in a cycle of DNA cutting and repair while the RNP complex remains present. NHEJ is therefore eventually error prone, and may lead to indels which can shift the reading frame of the protein encoded by that gene (5,6). This process functionally knocks out the encoded protein, which is therapeutically beneficial when a protein is overexpressed or has mutations through which the protein gained a pathogenic function. Therapeutically employing this mechanism is, for example, under clinical evaluation for the treatment of transthyretin amyloidosis through intravenous injection of LNP-formulated Cas9 mRNA and sgRNA (7). HDR, in contrast, causes partial resection of the broken DNA strand and uses a homologous DNA strand as template to guide the repair. This process naturally uses the sister chromatid during mitosis as the repair template. The exact mechanisms have been summarized excellently in other works (8–10). HDR can be exploited using an artificial DNA template to introduce specific mutations into a gene, and can therefore be used to repair damaged genes in genetic disorders. HDR has been used in this way to resolve point mutations as well as inserting larger DNA sequences (11–13).

However, HDR has proven to be difficult to translate to an effective gene therapy. HDR occurs less frequently than NHEJ, due to the relatively low expression of the effector proteins for HDR compared to NHEJ (8). Notably, HDR is active in the late S, G2 and early M phases of mitosis, but practically absent during other cell cycle phases (6,14,15). Furthermore NHEJ is always active and outcompetes the HDR machinery even during mitosis, leading to the odds of faithful gene correction to be low. This NHEJ preference by cells hampers clinical translatability of HDR, as the majority of treated cells will undergo the incorrect repair pathway and exhibit unwanted indels at the target site. Those cells are then no longer easy to target by CRISPR, as the target DNA sequence has mutated in an unpredictable manner and is now heterogeneous between edited cells. While autologous gene-corrected cells have recently entered clinical trials, this drawback has led the field to consider alternative gene-editing tools for direct *in vivo* injection of HDR machineries.

Prominent novel developments towards this include the Base-Editor and Prime-Editor systems. These cause single stranded DNA breaks and contain an additional effector protein domain fused to the Cas9 scaffold. Base editors chemically modify nucleic acids through their enzymes, whereas prime editor uses a reverse transcriptase and a modified guide RNA molecule to write mutations into the genome directly (16,17). The range of mutations these systems can resolve is in theory limited compared to HDR as these systems cannot facilitate large insertions, but preliminary results show that the specificity of the gene correction is much higher, especially for point mutations (17). These developments pose the question whether HDR mediated gene correction for small mutations is relevant for clinical development, possibly with add-on therapies to enhance its specificity.

Due to the aforementioned competition with NHEJ, it is essential that the repair pathway is shifted towards preferential or exclusive HDR before it can be safely used clinically. Many groups have demonstrated that small molecule compounds influencing the DNA repair pathways or the cell cycle are capable of improving HDR as recently reviewed by Shams et al. (10). Primarily efforts were done specifically on utilizing both NHEJ inhibitors and HDR enhancers. Prominent examples include SCR-7, a DNA ligase 4 inhibitor, which inhibits NHEJ and has been demonstrated to result in HDR becoming the dominant pathway both *in vitro* and *in vivo* (10,18,19). Direct HDR enhancement can be achieved by RS-1, which stabilizes the active conformation of Rad51, which is a limiting factor in HDR progression. This compound shows similar success as SCR-7 (10,20). Furthermore, alternative strategies utilizing HDR such as *in* trans Cas9 nickases can be utilized to improve the outcome of gene editing (21). While the *in vivo* data is promising, neither compound is in clinical development, making information on use in humans sparse.

The rationale of this work was therefore to explore a selection of clinically-tested drugs for potential CRISPR-modulating properties, to aid clinical development in future applications of HDR. We focused on drugs that are able to target DNA repair pathway regulation, signaling for cell proliferation and genomic instability in general (22). Interestingly, many therapies that are designed for cancer modulate proteins in these domains, as these proteins have an important role in both cancer and mechanisms involved in CRISPR gene editing. Therefore the aim was to screen oncological drugs to find novel modulators and pathways to enhance CRISPR HDR and enable potential add-on therapies in the future, and to add on to the growing toolkit of CRISPR enhancers used in the laboratory setting with clinically relevant drug molecules.

## Materials and methods

### HEK293T-eGFP and Hepa 1-6-eGFP cell culture conditions

HEK293T cells with constitutive enhanced green fluorescent protein (eGFP) expression (HEK293T-eGFP (23)) were cultured as described previously using low glucose DMEM containing 10% fetal bovine serum (Sigma-Aldrich, Zwijndrecht, The Netherlands) (12). Hepa 1-6-eGFP cells were cultured in high-glucose DMEM (Sigma-Aldrich) supplemented with 10% fetal bovine serum (Sigma-Aldrich). Cell culture plastics were acquired from Greiner Bio-One (Alphen aan de Rijn, The Netherlands).

Unless specified otherwise, gene editing experiments for both cell lines were conducted by seeding cells in 96-well Greiner CellStar plates (Greiner Bio-One) at a density of 3*10^5^ cells/cm^2^. The same cell density was applied in other well plate formats. Medium was supplemented with 1x antibiotic/antimycotic solution (Sigma-Aldrich) during gene-editing experiments, 48 hours after adding genome editing formulations.

### Hepa 1-6 eGFP cell line construction

Hepa 1-6 cells were graciously donated by dr. Piter Bosma from the Tytgat Institute for Liver and Intestinal Research, Amsterdam University Medical Centers. These cells were stably transduced using a lentiviral vector to constitutively express eGFP. Lentiviral particles carrying the eGFP gene were generated as reported previously (23). Briefly, lentivirus was made by co-transfection of a functional eGFP gene in the pHAGE2-EF1a-IRES-PuroR lentiviral vector, alongside the pMD2.G plasmid and PSPAX2 plasmid (Addgene #12259 and #12260, respectively) at a 2:1:1 ratio in HEK293T cells using 3 μg polyethylenimine (25 kDa linear, Polysciences, Warrington, USA) per μg plasmid DNA. The supernatant of these cells was cleared of cells by five minutes of centrifugation at 500 x *g*, followed by 0.45 μm syringe filter filtration. Lentiviral supernatants were stored at -80 °C until further use. Transduction was performed overnight at a multiplicity of infection of 0.1. Puromycin selection was performed using 2 μg/mL puromycin (InvivoGen, Toulouse, France) to the culture medium 48 hours post transduction. After two weeks of puromycin selection, eGFP expressing cells were sorted on a BD FACSAria III cell sorter (Becton Dickinson, New Jersey, USA), and subsequently expanded for 2 weeks prior to experimental use.

### Drug compound addition

A selection of forty small molecule drug compounds was made from the in-house oncological library of the high-throughput screening facility of the Princess Màxima Center. The selection was based on the mechanism of action of the drugs predicted to influence CRISPR outcomes. An overview of the compounds used in this study is given in Table 1. These compounds, dissolved in DMSO at 10 mM, were added to wells using the Echo 550 liquid handler (Beckman Coulter, Woerden, The Netherlands) for the large compound screen and the TECAN D300e digital dispenser (Tecan Group LTD, Männedorf, Switzerland) for the subsequent validation experiments. Cells were seeded in sterile cell culture plates pre-primed with the compounds calculated to yield the correct concentrations in each well. The concentration of DMSO was normalized in each well to 0.1% for all experiments and conditions in this work unless specified otherwise.

**Table 1:** Selected compounds and their respective IC50 in HEK293T-eGFP cells after 3 days of incubation. The dosage schemes in the gene-editing screening experiment were based on these IC50 values, see color coding. Compounds marked green had no measurable toxicity below 10 μM.

Legend and dosing scheme ( μM)
| High dose | Medium dose | Low dose |
| --- | --- | --- |
| 0.001 | 0.0001 | 0.00001 |
| 0.01 | 0.001 | 0.0001 |
| 0.1 | 0.01 | 0.001 |
| 1 | 0.1 | 0.01 |
| 10 | 1 | 0.1 |

| Compound | Target | IC50 (μM) |
| --- | --- | --- |
| Paclitaxel | TUBB | 0.00312 |
| Prexasertib | CHEK1 | 0.00533 |
| GSK461364 | PLK1 | 0.00835 |
| Romidepsin | HDAC 1 & -2 | 0.0164 |
| Volasertib | PLK1 | 0.0245 |
| Panobinostat | HDAC 1-11 | 0.0306 |
| THZ1 | CDK7 | 0.0371 |
| Adavosertib | WEE1 | 0.227 |
| CYC065 | CDK2 & -3 | 0.255 |
| Berzosertib | ATR | 0.335 |
| THZ531 | CDK12&13 | 0.352 |
| AT7519 | CDK1 & -2 | 0.438 |
| Karonudib | MTH1 | 0.450 |
| Birabresib | BRD2-4 | 0.540 |
| BI 894999 | BRD4 | 0.582 |
| CPI-203 | BRD4 | 0.651 |
| Abemaciclib | CDK4 & -6 | 1,07 |
| Belinostat | HDAC 1-11 | 1,10 |
| GSK1070916 | AURKB & -C | 1,39 |
| Molibresib | BRD4 | 1,78 |
| LMK-235 | HDAC4 & -5 | 1,79 |
| Ceralasertib | ATR | 1,84 |
| Vorinostat | HDAC 1-11 | 3,34 |
| Entinostat | HDAC 1-11 | 3,56 |
| Pevonedistat | NAE1 | 4,74 |
| I-BRD9 | BRD9 | 8,94 |
| Alisertib | AURKA | >10 |
| CPI-455 | pan-KDM5 | >10 |
| Epidaza | HDAC 1-3 | >10 |
| GSK2830371 | WIP1 | >10 |
| JQ-1 COOH | BRD4 | >10 |
| KU-55933 | ATM | >10 |
| KU-60019 | ATM | >10 |
| Olaparib | PARP1 & -2 | >10 |
| Pamiparib | PARP1 & -2 | >10 |
| PCI-34051 | HDAC 8 | >10 |
| Ribociclib | CDK4; CDK6 | >10 |
| Rucaparib | PARP1 & -2 | >10 |
| TAK-580 | BRAF; RAF1 | >10 |
| XAV-939 | TNKS1 & -2 | >10 |

### Cytotoxicity assays

Forty microliters of a HEK293t-EGFP cell suspension (3000 cells/well) were plated in tissue-culture treated flat-bottom 384-well microplates (catalogue number 3764, Corning, New York, USA) using a Multidrop Combi Reagent Dispenser (Thermo Scientific, Breda, The Netherlands). Cells were cultured for 16 to 24 hours under standard culturing conditions (5% CO_2_, 37 °C). Subsequently, 100 nL of the drugs (in DMSO, at different concentrations) are added to the wells containing the cells, to yield final concentrations of 0.1 nM, 1 nM, 10 nM, 100 nM, 1 μM and 10 μM (0.25% DMSO). All dose ranges were added in duplicate, followed by 72 hours of incubation. Cell viability was determined using a tetrazolium based metabolic activity assay (24). Briefly, 5 μL of 3-(4.5-dimethylthiazol-2-yl)-2,5-diphenyltetrazolium bromide (MTT) solution (5 mg/mL MTT in sterile PBS) was added per well, and the microplates were incubated for 4 hours at 37°C and 5% CO_2_ in a cell culture incubator. Next, 40 μL of 10% SDS/0.01 M HCl was added per well, and the microplates were incubated for 24-72 hours at 37°C and 5% CO_2_ in a cell culture incubator. Subsequently the absorbance at 570 nm and background absorbance at 720 nm were measured using the Spectramax i3x (Molecular Devices, San Jose, USA). Subsequently, the absorbance values at 720 nm were subtracted from the absorbance values at 570 nm, and the corresponding values were used to plot dose-response curves.

The data was normalized to the DMSO-treated cells (defined as 100% viability) and the empty controls (defined as 0% viability). IC50 values at 72 hours were calculated by determining the concentrations of the drug needed to achieve a 50% reduction in cell viability using the extension package *drc* in the statistic environment of R Studio (version 4.0.2) (25).

A narrower cytotoxicity range was determined by exposing cells to 0.1 - 1 μM of the tested compounds. 48 hours after treatment started, cells were washed and harvested using medium and one third of the volume was transferred to a fresh 96 well plate, analogous to how cells are treated during gene editing experiments. Subsequently, cells were cultured for another 3 to 5 days. In the case of 5 days, one third of the cells was again transferred into a fresh 96-well plate on day 3 to allow enough space for logarithmic cell growth during the additional incubation time. Cell viability was determined by the Promega One Step MTS assay (Promega, Madison, USA) using the manufacturer’s specifications. Relative cell viability for the Hepa 1-6-eGFP cells was calculated by normalizing the compound conditions to controls treated with DMSO only or DMSO plus gene editing formulations. For the HEK293T cells, the absolute absorbance at 590 nm was used as it was more representative of the relative cell viability between samples.

Cell morphology was assessed using the Nikon Eclipse Ti2 microscope (Nikon Europe, Amstelveen, The Netherlands). Pictures were acquired at 10x magnification with a Nikon DSLR 10 camera using the same imaging settings within each experiment set (Nikon Europe, Amstelveen, The Netherlands).

### CRISPR-Cas9 nanocarrier formulation

SpCas9 protein was produced and purified in-house as described previously (12). sgRNA and HDR template DNA were acquired from Sigma-Aldrich (Haverhill, United Kingdom).

Lipid nanoparticles (LNP) carrying SpCas9, sgRNA and HDR template DNA were formulated using the components and molecular ratios described previously (12). 1,1′-((2-(4-(2-((2-(bis(2-hydroxydodecyl)amino)ethyl)(2-hydroxydodecyl)amino)ethyl)piperazin-1-yl)ethyl)azanediyl)bis(dodecan-2-ol) (C12-200) was acquired from CordonPharma (Plankstadt, Germany) and used as the ionizable lipid in the formulation. 1,2-dioleoyl-sn-glycero-3-phosphoethanolamine (DOPE) was acquired from Lipoid GmbH (Steinhausen, Switzerland), Cholesterol and 1,2-dimyristoyl-rac-glycero-3-methoxypolyethylene glycol-2000 (PEG-DMG) were acquired from Sigma-Aldrich, and 1,2-dioleoyl-3-trimethylammonium-propane (DOTAP) was acquired from Merck (Darmstadt, Germany). LNP were produced using microfluidic mixing with the Dolomite Microfluidics system (Dolomite Microfluidics, Royston, United Kingdom) and herring-bone micromixer chip with hydrophilic coating (Dolomite Microfluidics, catalogue number 3200401). A total flow rate of 1.5 mL/min and flow rate ratio of 2:1 were used between an aqueous outer phase containing SpCas9, sgRNA and HDR template in nuclease free water, and the lipids in 100% ethanol in the inner phase, respectively. The resulting LNP were diluted 4 times in Dulbecco’s PBS (Sigma-Aldrich). In the experiments using the Hepa 1-6 eGFP cells, ProDeliverIN CRISPR (OZ Biosciences, San Diego, USA) was used as reported previously (12).

### Compound screening to modulate CRISPR repair outcomes

Compounds were assessed in three dosages to assess the effects on gene-editing efficacy. The highest concentration was based on the IC50 of the compounds as determined by the cytotoxicity determination, with a medium and low dose which were 10 and 100 times diluted respectively compared to this highest dosage. Cells were incubated with these compounds for 24 hours prior to LNP addition. LNP were added to all wells at a final concentration of 20 nM SpCas9 to achieve robust genome editing (12). After 24 hours of co-incubation, medium was refreshed and cells were transferred to a 48 well plate for further culturing. Six days after adding compounds, cells were processed for flow cytometric analysis of the gene knock-out and gene-correction efficiencies (26). Briefly, a ssDNA template was used carrying two nucleotide mutations to convert the eGFP sequence to that of a blue fluorescent protein (BFP), as well as mutating the PAM sequence to ensure robust HDR. The sgRNA spacer sequence was 5’-GCUGAAGCACUGCACGCCGU-3’, and the HDR template sequence was 5’-CAAGCTGCCCGTGCCCTGGCCCACCCTCGTGACCACCCTGAGCCACGGCGTGCAGTGCTTCAGCCGCTACCCCGACCACATGAAGC-3’.

Flow cytometry using the BD FACS Canto II (Becton Dickinson) was used to determine cells undergoing NHEJ (eGFP and BFP negative population) and HDR (eGFP negative, BFP positive). Data analysis was performed using Flowlogic (version 8.7). Graphpad PRISM (version 9.3.1) was used for statistical analysis and preparing graphs.

The percentage of HDR relative to total gene editing in a given cell population (hereafter named Relative HDR (% of edited cells) was calculated by dividing the absolute HDR population by the sum of gene-edited cells found in the BFP+ and eGFP-gates. Absolute HDR (% of all cells) and Absolute NHEJ (% of all cells) were analyzed where appropriate. The gating strategy is given in Supplementary Figure 1.

### Gene sequencing and genotype analysis

For genotypic analysis, HEK293T-eGFP cells were treated with alisertib for 72 hours and CRISPR LNP for 48 hours prior to harvesting by trypsinization. 25% of the harvested cells were transferred to a fresh well plate for expansion and analysis by flow cytometry as described previously. The remaining cells were lysed and genomic DNA was extracted using the QIAamp DNA Blood Mini Kit (Qiagen, Venlo, The Netherlands) according to the manufacturer’s instructions. PCR was performed to amplify the eGFP locus in the obtained DNA using the Phusion™ High-Fidelity DNA Polymerase (2 U/μL) (Thermo Fisher Scientific, Landsmeer, The Netherlands). The PCR mixture (50 μL) contained 200 ng of DNA, 0.5 μM of forward primer (5’-GACGTAAACGGCCACAAGTT - 3’ (Integrated DNA Technologies Leuven, Belgium) and reverse primer (5’-CGATGTTGTGGCGGATCTTG - 3’ (Integrated DNA Technologies Leuven, Belgium), 200 μM of dNTPs (dNTP Mix (10 mM each) (Thermo Fisher Scientific), 1 × Phusion HF buffer, 3% DMSO and 1 units of Phusion High Fidelity DNA polymerase. The DNA was amplified using the following thermocycling steps: 98°C for 30 sec; 35 cycles of 98°C for 10 sec, 62°C for 30 sec and 72°C for 30 sec; 72°C for 10 min. The PCR products were purified using the GeneJET PCR Purification Kit (Thermo Fisher Scientific). Sanger sequencing was performed by Macrogen (Amsterdam, The Netherlands) using the previously mentioned reverse primer as sequencing primer. Sanger sequencing chromatograms were analyzed using the TIDER webtool (http://shinyapps.datacurators.nl/tider/) using default settings (27). The reference chromatogram, corresponding to the blue fluorescent mutation, was generated from a gBlock gene fragment acquired from Integrated DNA technologies. The control (eGFP) chromatogram was generated from untreated cells. The indel frequencies up to -5 and +5 were plotted using Graphpad PRISM, version 9.3.1.

## Results

Compounds were selected from a clinically assessed oncological drug library used for drug discovery in pediatric cancer, as explained in figure 1A. This library contained a large variety of drugs, prompting a selection to be made. The rational drug selection was done by defining groups of pathways expected to modulate CRISPR gene editing outcomes: cell cycle modulation, DNA damage repair modulation and chromatin modulation. Drugs which were not easily categorizable in a single domain were included in the screen as well, due to the potential of unexpected effects on the gene editing outcome. This selection led to 40 compounds to be screened, summarized in Table 1.

**Figure 1.**
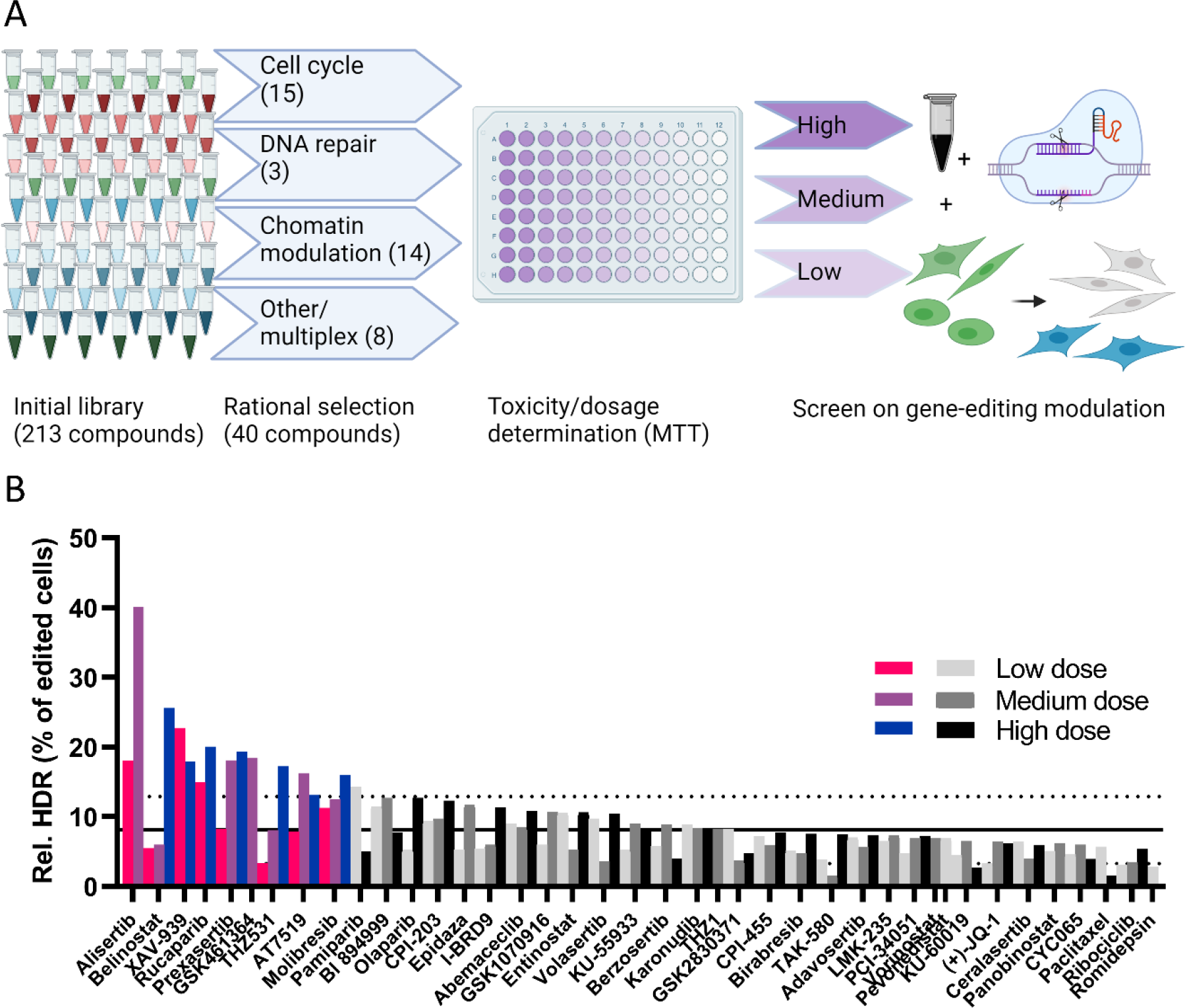
Initial screening performed using oncological compounds on CRISPR genome editing outcomes. A: Scheme showing the selection and screening process of the compounds for toxicity and gene-editing efficiency evaluation. Toxicity screening using a cell viability assay was done to find the dosages to be used in the subsequent drug screening for effects on gene-editing efficiency by conversion of eGFP positive cells to nonfluorescent or blue fluorescent cells. B: Efficiency of HDR relative to all gene edited cells, in HEK293T-eGFP cells treated with compounds in up to three dosages based on toxicity screening: High (^10^log value below IC50), Medium (10-fold lower than high dose) and Low (10-fold lower than medium dose). Efficacy was compared to cells treated with only CRISPR LNP (mean +-SD as solid and dotted lines; n=29 wells). Each bar represents one well of >1000 cells in the single-cell gate in flow cytometry. Colored bars were considered hits in this initial screening, and studied in further validation experiments (explained in text).

The cytotoxicity of the selected compounds was first assessed on the HEK293T-eGFP cell line using an MTT assay and determining the IC50 values of each compound after three days of treatment. These conditions were selected to mimic the maximum exposure time to the compound to be used in the screen (Table 1, Supplementary Figure 2). Three compounds showed considerable toxicity with IC50 values in the nanomolar range. The majority of compounds were tolerated in the micromolar range however, or were not toxic in the investigated concentration range. These toxicity values were used to dose the compounds in a sub-toxic dosage in subsequent gene-editing experiments, as noted in Table 1. The closest ^10^log concentration to the IC50 (high dose) and two ^10^log values below were used to preliminarily determine effects of the compounds on gene-editing efficiency, as the compound needed to be efficacious in a non-toxic or at most subtoxic dose for potential therapeutic application. Cells were incubated with the compounds for 24 hours prior to adding lipid nanoparticles (LNP) carrying Cas9 enzyme, sgRNA and an HDR template designed to mutate the eGFP sequence to a blue fluorescent phenotype, as reported previously (12).

**Figure 2.**
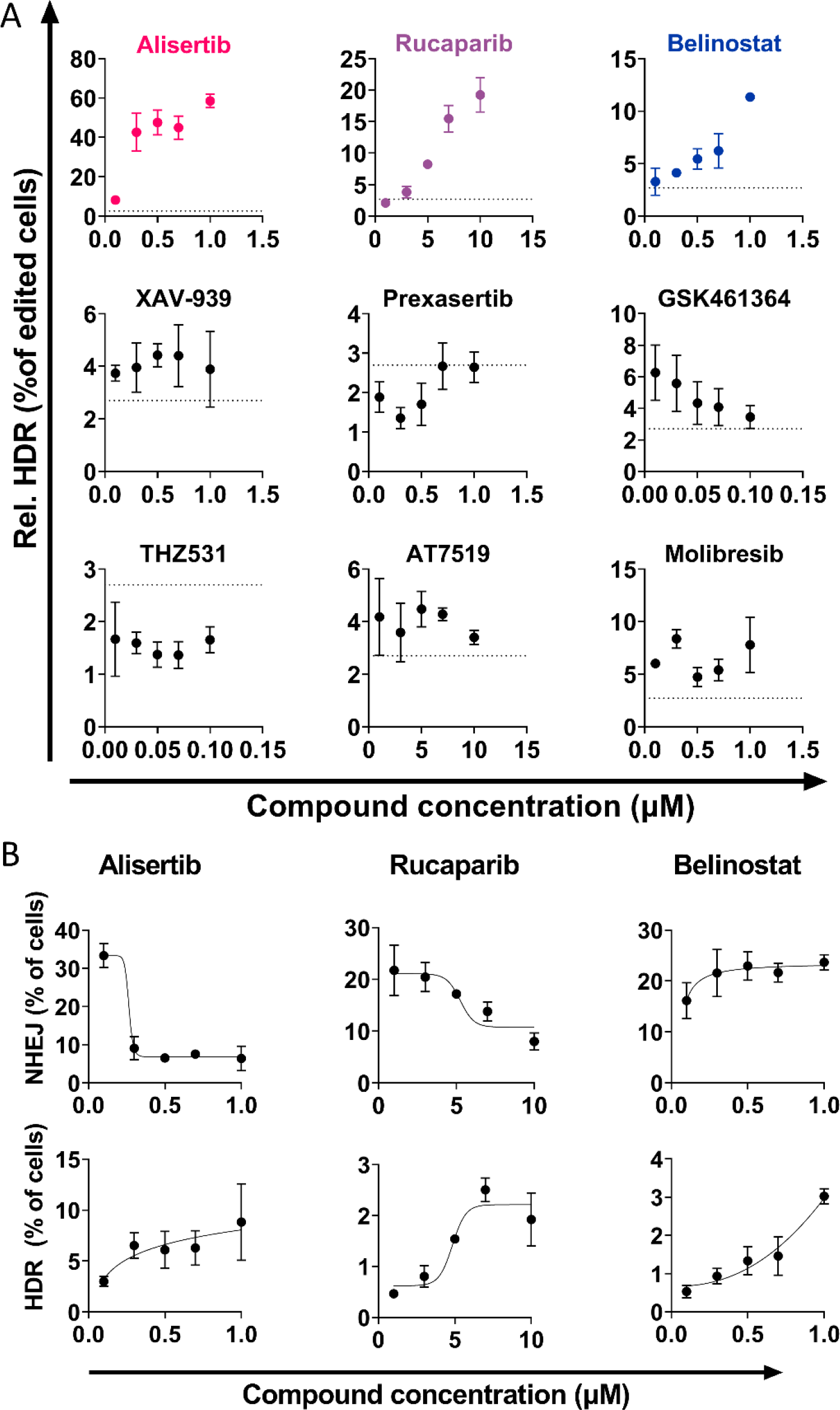
Hit validation of the findings in Figure 1. A: Repeated experiment in a narrower dose-range for the hits compared to LNP treatment without compounds (mean (dotted line); 2,9%). Of these, alisertib, rucaparib and belinostat yielded a clear dose-dependent HDR increase. B: The result of the three significant and dose-dependent hits from A separated into the gene knock-out (NHEJ, top) and correction (HDR, bottom) outcomes.

The effect of all screened compounds on gene editing are given in figure 1B. Gene knockout (loss of eGFP signal) and correction (rise of BFP signal) were measured, as shown in Supplementary Figure 3. From these values the relative HDR efficiency was calculated as the percentage of HDR in total gene edited cells, as shown in Figure 1B. The gating strategy used in the flow cytometry data analysis is given in Supplementary Figure 1.

**Figure 3.**
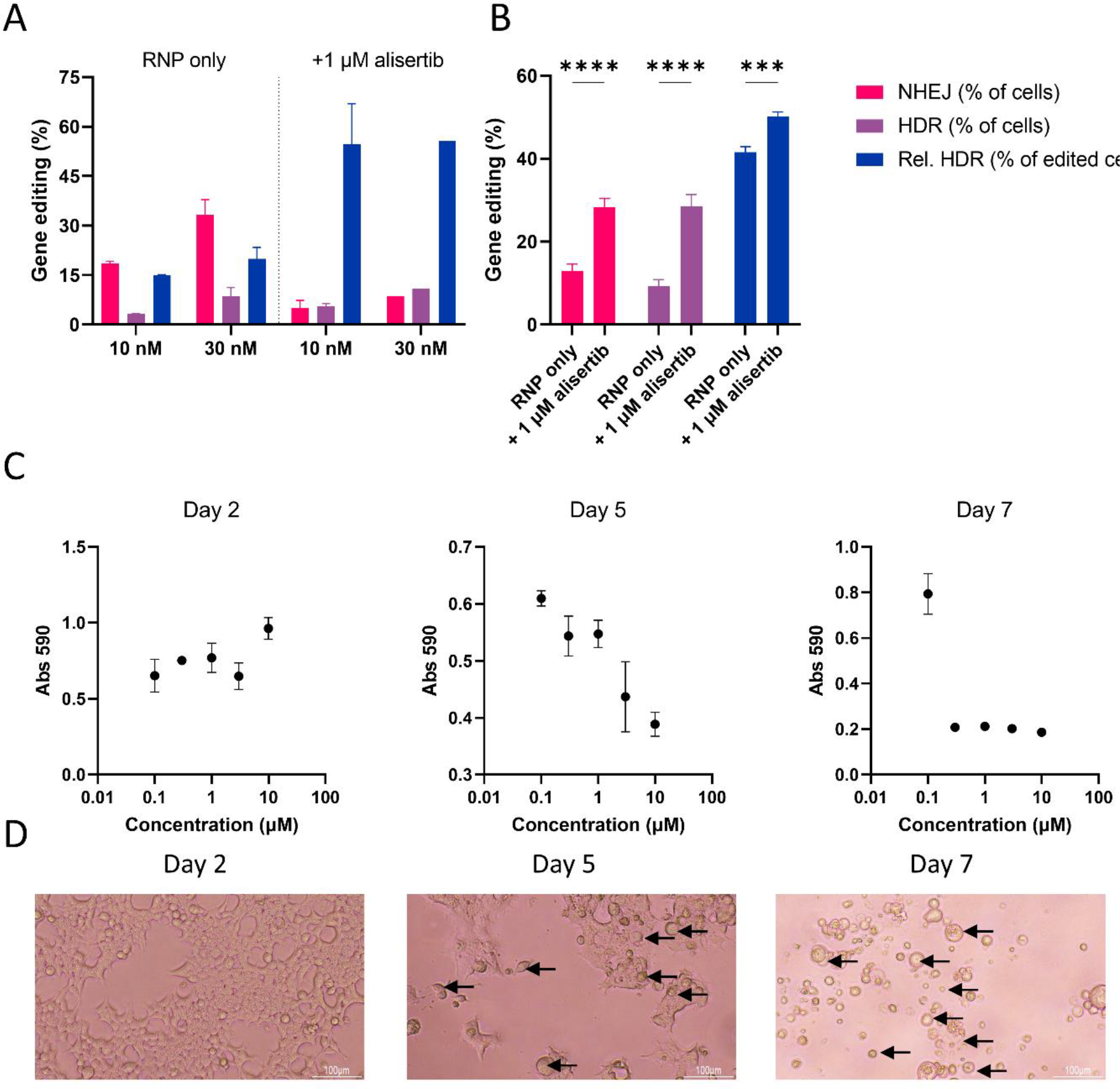
Further validation of alisertib in HEK293T cells relating to CRISPR gene editing outcomes. A: LNP dose-dependency in cells treated without or with 1 μM alisertib. B: TIDER gene-editing outcomes for cells treated with 10 nM CRISPR formulation alone or co-treated with 1 μM alisertib. C: Time-resolved toxicity of cells treated with alisertib at 0 days. Medium containing alisertib was replaced with standard culture medium on day 2.

The effect of the compounds was compared to cells treated with only LNP containing RNP and HDR template DNA (n=29 wells). A minimum of 1000 events in the single-cell gate was deemed necessary at a minimum for data analysis. Conditions not exceeding this number due to unexpected toxicity were excluded from figure 1B. Compound treated cells deviating at least one standard deviation from the LNP-only control mean (dashed line in figure 1B) were considered to differ relevantly from LNP alone, and were considered a potential hit for altering the gene-editing outcome selection. In this study, only compounds which increased the relative incidence of HDR were investigated further, other findings are summarized in Supplementary Figure 2. This was calculated by the amount of HDR events divided by all gene edited cells (blue fluorescent and non-fluorescent cells combined). Nine compounds showed at least one concentration above that threshold. Further conclusions were not taken from this screen as all datapoints were single measurements. Validation experiments were performed to confirm these hits in triplicate and in a narrower dosage range.

The hits were validated by narrowing the dose range between the most efficacious concentration found in the initial screen and the ^10^log-lower concentration in triplicate (Figure 2A). Three compounds showed a strong dose-dependent preferential activation of HDR over NHEJ compared to controls treated with LNP only (dotted line): rucaparib, belinostat and alisertib. The other compounds did not show a clear dose-dependent enhancement of HDR efficiency upon this further scrutiny. Further analysis was done on the two main gene editing outcomes of NHEJ and HDR. The three validated hits exhibited different effects on these two repair pathways as shown in Figure 2B. Rucaparib and alisertib both inhibited NHEJ and improved HDR, while belinostat increased both NHEJ and HDR, with a relatively pronounced increase for HDR in this study. Taken together, alisertib exhibited the strongest effect on both gene editing outcomes (NHEJ inhibition and HDR enhancement) compared to LNP-treated control cells. Between 0.1 and 0.3 μM the relative HDR incidence increased over 5-fold and became the preferred gene editing outcome (>50% relative HDR incidence) at 1 μM. Due to this drastic effect, a narrower dose-range was investigated, shown in Supplementary Figure 4. The inhibitory effect on NHEJ was dose-dependent in this range, while HDR markedly increased between 0.2 and 0.3 μM. Thus 0.3 μM seemed to be the lowest effective concentration for preferential HDR activation in HEK293T-eGFP cells, while 1 μM caused HDR to become the most prominent repair pathway.

**Figure 4.**
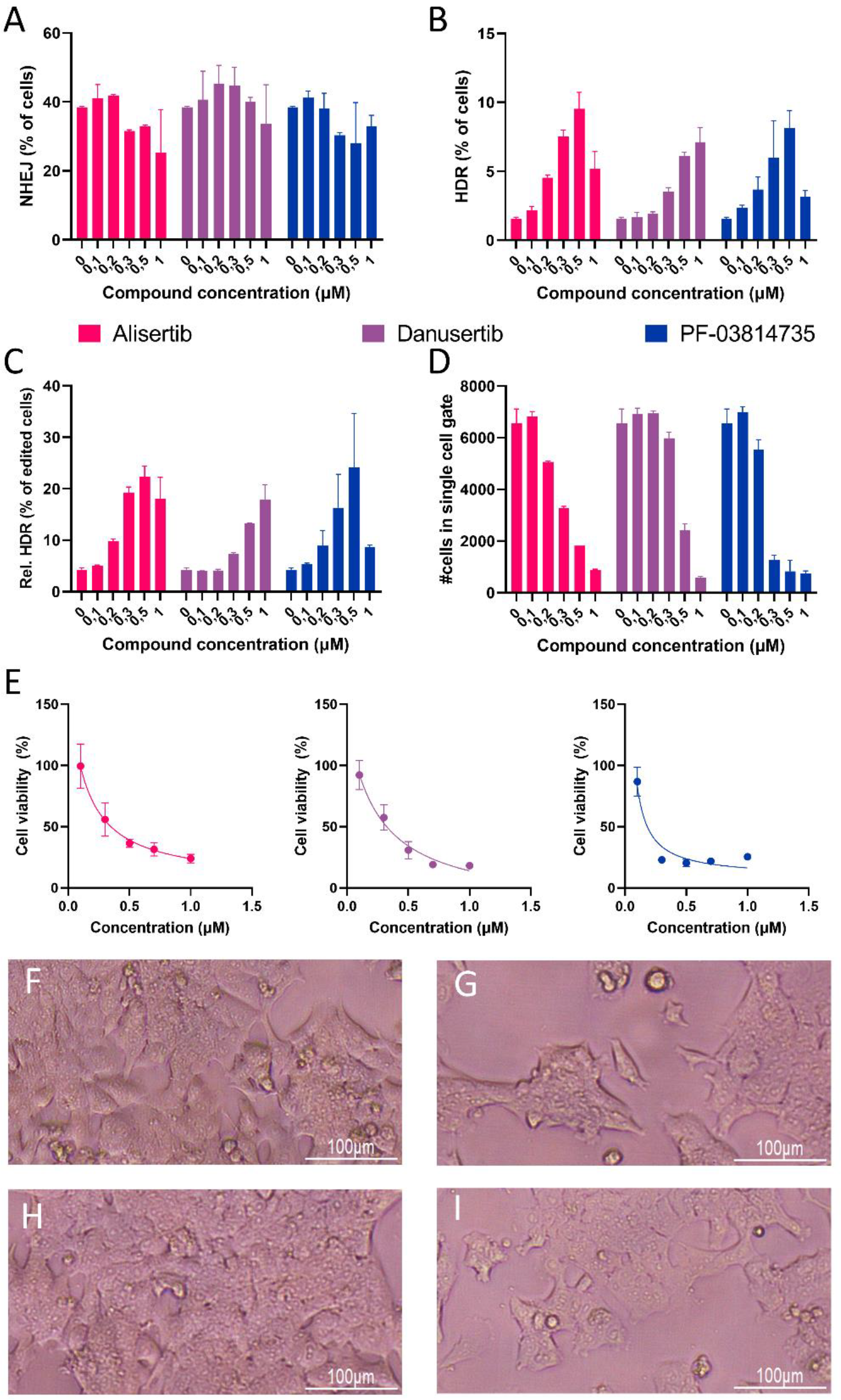
Gene editing efficacies and cytotoxicity on Hepa 1-6 eGFP pretreated with aurora kinase inhibitors alisertib, danusertib or PF-03814735 using ProdeliverIN CRISPR for RNP delivery. Colors are consistent between panels A-E. A: NHEJ incidence with ascending dosages of the three AURKA inhibitors. B: Absolute HDR incidence with ascending dosages of the thee AURKA inhibitors. C: Relative HDR incidence calculated from the percentages in panels A and B, for cells treated with ascending dosages of the three AURKA inhibitors. D: Cell counts in the single cell gate in the flow cytometry data after acquiring 10.000 cells per condition, for cells treated with ascending dosages of the three AURKA inhibitors. E: Cell viability measured by MTS assay 5 days after the start of treatment. F-I: Microscopic pictures of cells treated with no compound (F), 0.3 μM of alisertib (G), 0.3 μM danusertib (H) or 0.3 μM PF-03814735.

To assess the dose-dependency of alisertib on the CRISPR-Cas mediated gene editing outcome, cells were pre-treated with either 0 or 1 μM alisertib 24 hours prior to LNP addition. LNPs were administered to cells with either 10 nM (standard dosage in other experiments) or 30 nM of SpCas9. When a higher dose of LNP was added, both gene knock-out and gene correction populations increased proportionally to the dosage as shown in figure 3A. In the case of pre-treatment with 1 μM alisertib, the relative HDR incidence stayed above 50% indicating that with a higher total gene editing incidence, HDR was still the predominant pathway. Furthermore, when the alisertib incubation time was varied it showed that simultaneous treatment improved gene editing, with a 2.5-fold significant increase of relative HDR efficiency (Supplementary Figure 5).

HDR-mediated gene correction was further validated at the genetic level by amplifying the eGFP locus using PCR and subsequent sequencing of the amplicons. The sequencing traces were analyzed using the TIDER method (27). This showed that at the genetic level, the relative HDR incidence was higher for alisertib primed cells as well. However, the total NHEJ and HDR incidences found by TIDER analysis were much higher than found in the fluorescent protein expression in flow cytometry. The distribution of insertions and deletions revealed that most genotypes had a -3 deletion, which could explain this discrepancy as this may not lead to gene knockout in some cases (Supplementary Figure 6).

An observation in these experiments was that cells treated with alisertib had a delayed cytotoxicity, which was not captured in the initial MTT assay. After 2 days, the cell viability as measured by MTS was not affected by alisertib. This is shown in Figure 3C as well as Figure 3D, in which the morphology is shown to resemble healthy HEK293T cells. However after 5 days, consistent with the duration of the experiments presented in this work, cells started exhibiting a dose-dependent decrease in cell viability. The confluency decreased, and cells with a disturbed morphology started appearing (Figure 3D, marked by the arrows). Seven days after treatment started, cells treated with at least 0.3 μM alisertib showed very low metabolic activity and confluency, and cell morphology was completely disrupted.

Hepa 1-6-eGFP cells, a murine hepatoma cell line, were finally used to investigate whether the observed HDR preference was cell-line and species independent. Three Aurora kinase inhibitors were used with differing specificities for aurora kinases A, B and C, to assess the pathway specificity in parallel. Alisertib is selectively an AURKA inhibitor. PF-03814735 inhibits both AURKA and Aurora kinase B (AURKB) and danusertib is a pan-aurora kinase inhibitor of AURKA, AURKB and Aurora kinase C (AURKC). All three inhibited NHEJ up to 2-fold (Figure 4A) increased HDR up to 5 fold (Figure 4B). This resulted in a positive trend for improving relative HDR incidence similarly to HEK293T-eGFP cells, as shown in figure 4C. Relative HDR increased 3-fold for alisertib in concentrations higher than 0.3 μM, and similar effects were seen for danusertib and PF-03814735. However, toxicity was a concern in these cells as well. The number of detected cells in flow cytometry decreased with higher dosages (Figure 4D), which was due to a reduced cell viability as measured by metabolic activity (Figure 4E). A dosage of 0.3 μM relatively showed overall high efficacy and manageable toxicity for alisertib and danusertib, while 0.2 μM was favorable for PF-03814735. Taken together alisertib had a strong effect (22.4% relative HDR incidence) for a relative cell viability of 37% at a concentration of 0.3 μM, which is the most favorable profile between the three inhibitors and the tested concentrations. Microscopy revealed that the cell morphology after treatment with 0.3 μM danusertib (Figure 4H) after 5 days was not disrupted compared to untreated control conditions (Figure 4F). The morphology using alisertib (Figure 4G) or PF-03814735(Figure 4I) also did not change as drastically as it did for the HEK293T-eGFP cells, but the confluency of cells was noticeably lower than in the untreated control.

## Discussion

The initial rationale of this screen was to find compounds that lead to HDR being favored over NHEJ, which can feasibly be given as a targeted, synergistic treatment with CRISPR-Cas9-based gene editing therapeutics. Oncological drugs were screened due to the similarities between the pathways targeted in cancer and those involved in genome editing, such as cell cycle regulation and DNA damage repair. The 40 selected compounds exhibit varied subcellular targets and processes as shown in Table 1. Many studies on small molecule CRISPR enhancers have been performed already, with varying success. For example, the DNA ligase 4 inhibitor SRC7 has been widely utilized (18,19). However, this compound has not been used in any clinical trials, while many of the compounds investigated in this study are, or were, in various phases of clinical development.

The screen revealed many compounds that did not affect the outcome of gene editing significantly or relevantly, but also three that did show a favorable effect. Two out of three confirmed hits were reported to influence gene repair outcomes in previous studies. HDAC inhibitors, such as belinostat, have shown in the past to improve overall gene editing (28) and HDR specifically (29), due to their effects on chromatin packaging of the DNA. This efficacy was recently demonstrated for prime editing as well (30). These compounds therefore served as an internal validation for the screen. Interestingly however, the other pan-HDAC inhibitors (entinostat, vorinostat and panobinostat) did not show the same effect. Other HDAC inhibitors were not reported previously. Epidaza, which inhibits HDAC 1-3, did not show an effect towards improving HDR and romidepsin, which inhibits HDAC 1 and 2, strongly inhibited HDR in this study compared to NHEJ. Further study on which HDAC subtypes inhibited by these compounds dictate genome editing outcomes is therefore needed.

Rucaparib, a PARP 1 and PARP 2 inhibitor involved in the DNA damage signaling checkpoint, affected the gene editing outcomes as well. Whereas rucaparib has previously been shown to improve gene editing due to inhibition of the microhomology-mediated end joining pathway (31), the observed effect on HDR found in the current study has not been reported to the best of our knowledge. Inhibition of PARP 1 and PARP 2 directly influences the regulation of base excision repair, which is usually a single stranded DNA damage event. However it is reported that inhibition of PARP-1 drives the cell towards homologous recombination, which in oncology is used to cause cell death in BRCA-deficient cells (32), and could explain our observations. Furthermore, this drug is approved for clinical use in humans, and therefore clinical knowledge exists to potentially devise a synergistic treatment plan for CRISPR and rucaparib combination therapy.

The primary discovery of the screening was the simultaneous NHEJ inhibition and HDR induction found when pretreating cells with alisertib. This compound is used in anti-cancer therapy to inhibit AURKA, which is involved in mitotic spindle formation and organization, and has been implicated in DNA signaling in cancers (33,34). Reports on mechanisms in healthy cells are sparse, but the toxicity was shown to be lower in healthy cells than in breast cancer cells (35). The effects found on gene editing efficiency therefore needs to be investigated more on the mechanistic level to unravel this observed relationship between AURKA inhibition and HDR efficiency.

Addition of alisertib resulted in the greatest effect observed in this study, showing a preference for HDR over NHEJ outcomes in HEK293T-eGFP cells. This was seen on the phenotypic level by BFP expression compared to eGFP knock-out, and to a lesser extent on the genetic level shown by sequencing and TIDER analysis. This may be due to the predominant mutations found in TIDER being in-frame (−3), which may not disrupt the eGFP protein function. We found that treating cells with alisertib and treating them with a higher dosage of LNP increases the efficacy as well, validating further that the effect is due to priming the cells for CRISPR HDR by increasing the RNP and HDR template concentrations. If this pathway can be inhibited in a non-toxic way, it can therefore lead to greater specificity of gene correction.

In our initial screen we classified alisertib to be non-toxic, based on the IC50 gathered from the MTT assay data. However, when looking closer at the toxicity curve two days after treatment, a loss of 20% cell viability can be seen at a concentration of 0.1 μM (Supplementary Figure 2). This led us to scrutinize the toxicity in more detail. The toxicity becomes apparent 5 days after the start of alisertib treatment. This was independent of total compound incubation time, which was varied between 24 hours and 0 hours of pre-incubation of cells with alisertib (Supplementary Figure 5). The observed in the Hepa 1-6 cells after 5 days presented in figure 3 are in line with the HEK293T results, which indicates that the toxicity was simultaneously species and cell type independent, at least in these model systems. Toxicity of these AURKA inhibitors needs to therefore be investigated further in more relevant cell types to assess if these effects are transient and significant, as it might be possible that healthy cell types, rather than cancer cells, are more resistant to these compounds.

Finally, two other AURKA inhibitors (danusertib and PF-03814735) were assessed in Hepa 1-6-eGFP as well to validate the pathway. The manufacturer summarized the efficacy of these compounds towards AURKA, AURKB and AURKC. Of these, PF-03814735 is the most potent towards AURKA with an IC50 of 0.8 nM. Alisertib has potency in the same order of magnitude with an IC50 of 1.2 nM, and danusertib is magnitude less active at 13 nM. This is reflected in the efficacy, as danusertib requires a higher concentration before the effect on gene editing efficiency, as well as the toxicity, was visible, although the toxicity is in the same order of magnitude for all three compounds. Danusertib also has activity against AURKB and AURKC, with an IC50 of 79 and 61 respectively, whereas PF-03814735 has a preference for AURKB at an IC50 of 0.5 nM. GSK1070916, an AURKB and AURKC inhibitor, did not show an effect toward relative HDR activation, so these pathways likely only contribute to the cytotoxicity. PF-03814735 showed a clearly more drastic toxicity, likely due to the strong AURKA and AURKB inhibition. Danusertib and alisertib showed similar cytotoxicity, but the efficacy of alisertib was higher.

## Conclusion

Of the forty screened compounds, three showed a significant HDR enhancing effect: belinostat, rucaparib and alisertib. Alisertib specifically shows a rapid onset of action to this end, as well as activity in a relevant cell line. While AURKA inhibition showed a relevant increase of HDR, the toxicity displayed in this study limits its application. Other means of AURKA inhibition might be effective and warrants further investigation. Furthermore, targets downstream of AURKA should be further investigated to find the specific drivers of this effect, and allow application in an HDR-based gene editing approach in a more relevant setting such as primary cells or *in vivo*.

## Conflict of interest

The authors have no conflicts of interest to declare.

## Acknowledgements

The authors would like to acknowledge Omnia Elsharkasy for her help with cell sorting to select high-expressing Hepa 1-6 eGFP cells.

This research was funded by the Netherlands Organisation for Scientific Research (NWO) Talent program VICI, grant number 865.17.005.

## Contribution statement

**Danny Wilbie**: Conceptualization, Investigation, Methodology, Writing – Original Draft. **Selma Eising**: Conceptualization, Methodology, Formal Analysis. **Johanna Walther**: Methodology. **Vicky Amo-Addae**: Methodology, Investigation. **Esmeralda Bosman:** Methodology, Investigation. **Olivier Gerrit de Jong:** Resources, Writing – Review & Editing. **Jan Molenaar:** Conceptualization, Resources, Supervision. **Enrico Mastrobattista**: Conceptualization, Supervision, Writing – Review & Editing.

## Notes

### Competing Interest Statement

The authors have declared no competing interest.

