## Supplemental Information for "Anti-cancer compound screening identifies Aurora Kinase A inhibition as a means to favor CRISPR/Cas9 gene correction over knock-out"

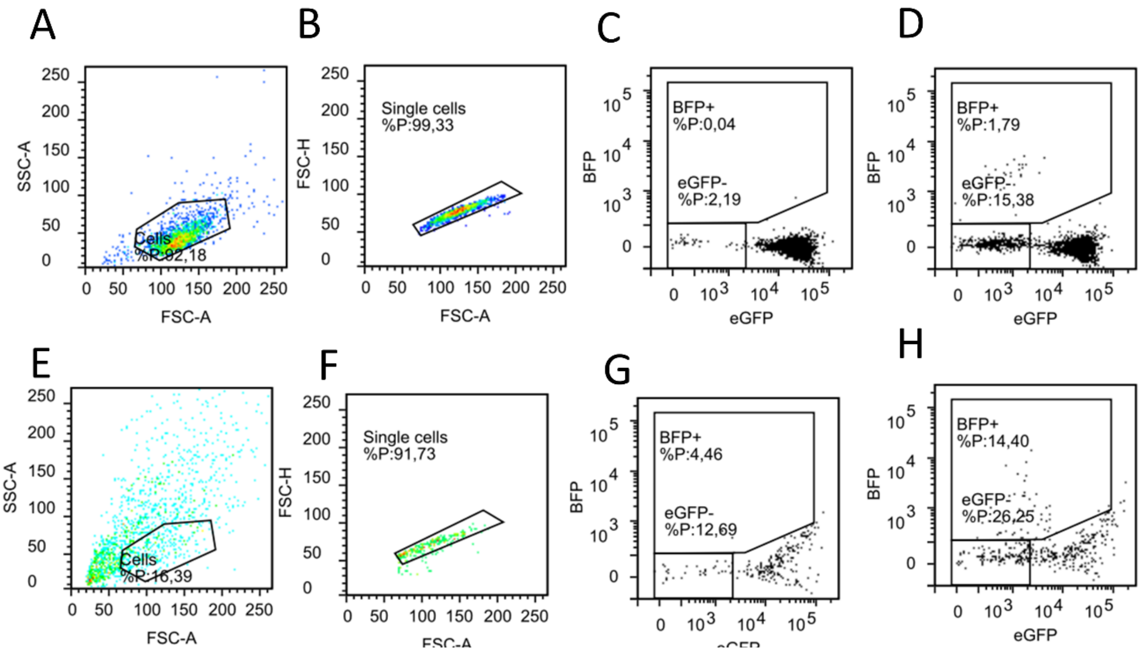


Supplementary figure 1: Gating strategy employed in typical flow cytometry measurements for gene editing outcomes in HEK293T-eGFP cells. A-D: Gating without additional compounds. E-H: Gating in presence of 1 µM alisertib. A and E: gating cells. In presence of alisertib the spread of forward and side scatter changes, it is assumed that cells with normal morphology, as seen in figure 3E, are in the same location in this dot plot. B and F: single cell gating. C and G: eGFP knock-out and BFP emergence in absence of gene editing LNP. D and H: eGFP knock-out and BFP emergence in presence of gene editing LNP.

NHEJ incidence was calculated in the eGFP$^-$ gate in D or H and subtracting the eGFP$^-$ gate from C or G respectively. Absolute HDR incidence was calculated in the BFP+ gate in D or H and subtracting the BFP$^+$ gate from C or G, respectively. Relative HDR incidence was calculated by dividing the absolute HDR incidence by the sum of NHEJ incidence and absolute HDR incidence.

Notable, alisertib treated cells had a high “false positive” rate in the eGFP$^-$ and HDR gates in the control (G), which were subtracted from relevant results.

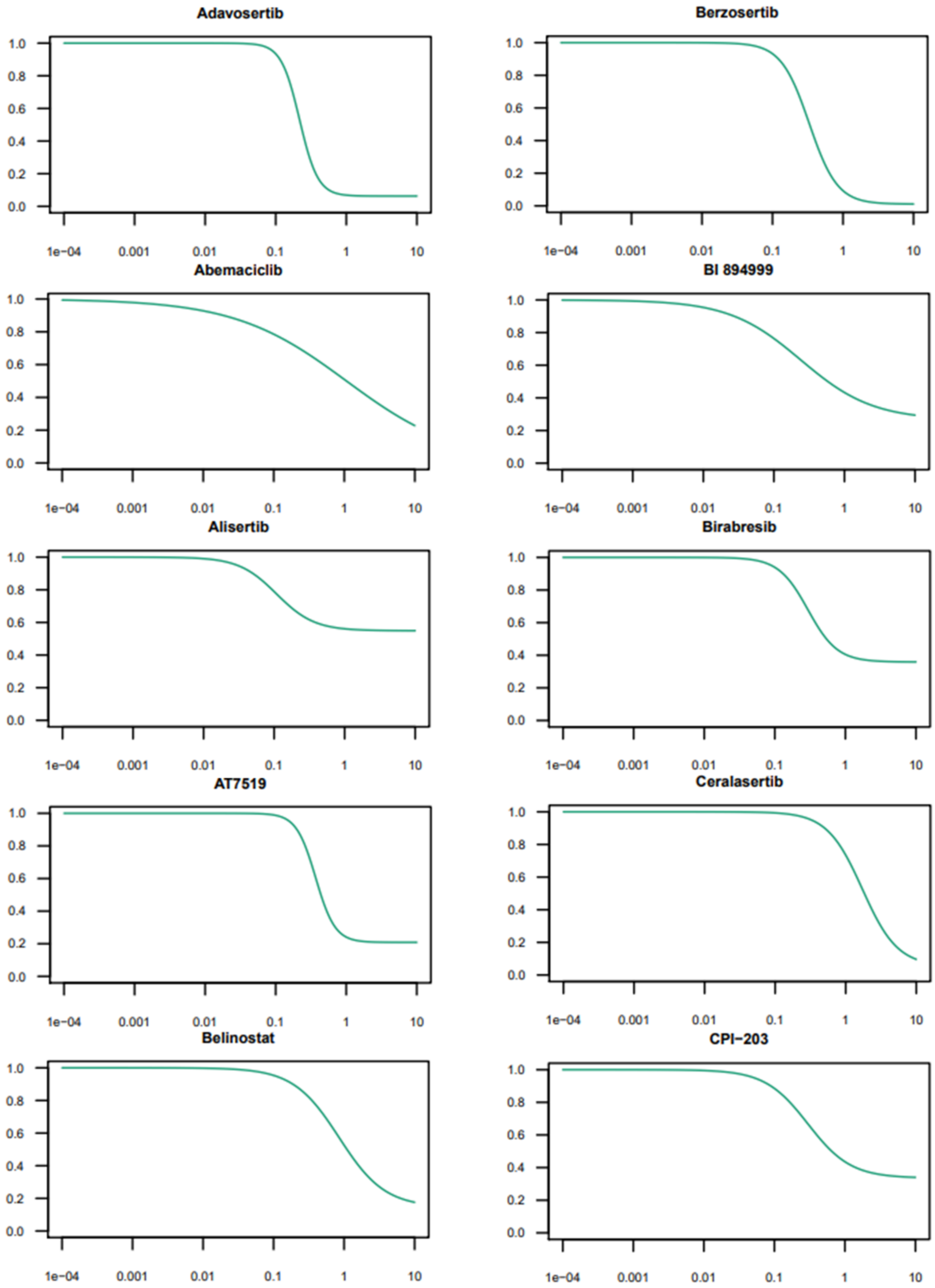
Adavosertib
Berzosertib
Abemaciclib
BI 894999
Alisertib
Birabresib
AT7519
Ceralasertib
Belinostat
CPI−203
1.0 0.8 0.6 0.4 0.2 0.0
1e−04 0.001 0.01 0.1 1 10

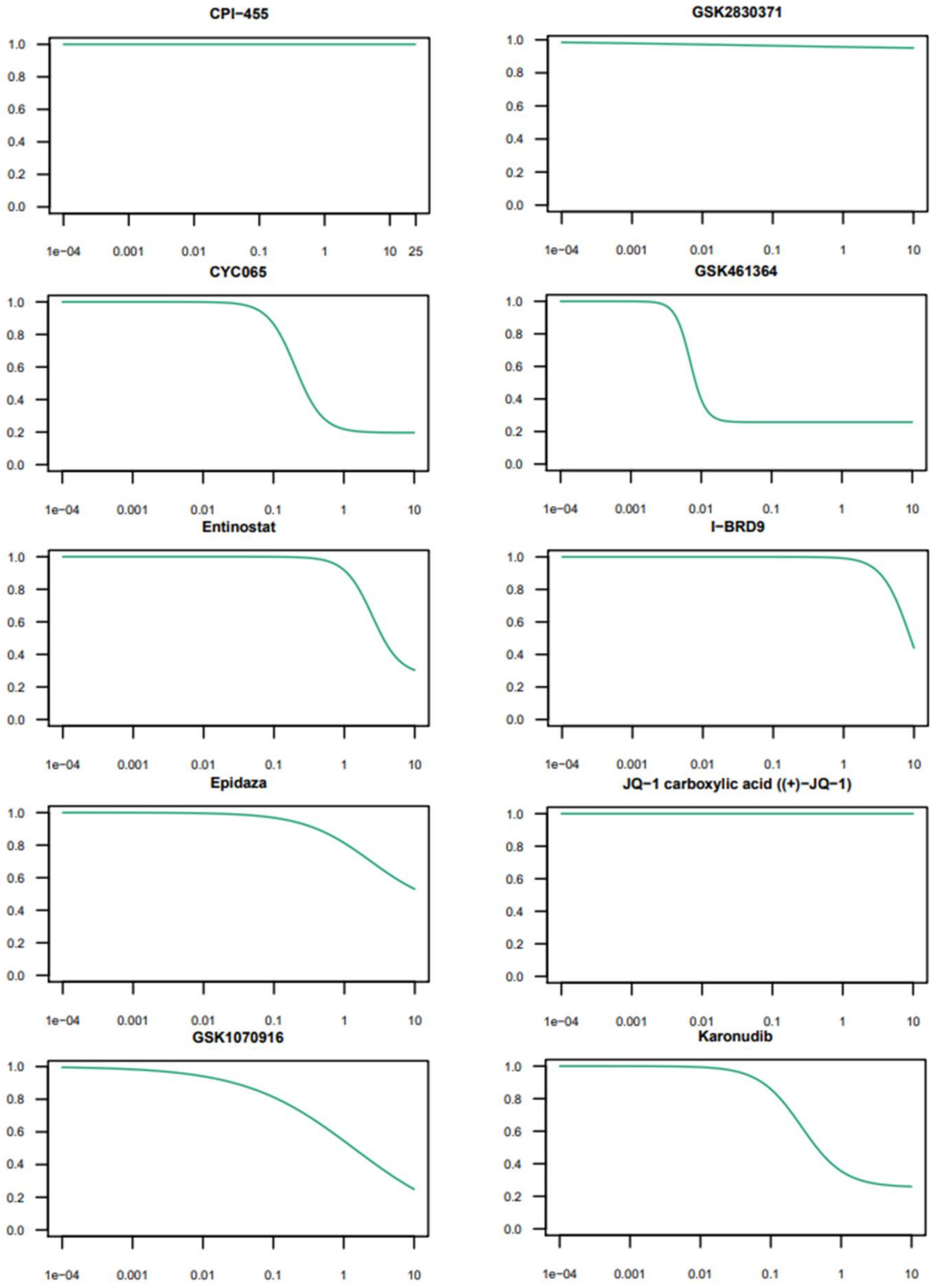
CPI−455
GSK2830371
CYC065
GSK461364
Entinostat
I−BRD9
Epidaza
JQ−1 carboxylic acid ((+)−JQ−1)
GSK1070916
Karonudib
0.0 0.2 0.4 0.6 0.8 1.0
1e−04 0.001 0.01 0.1 1 10 25
1e−04 0.001 0.01 0.1 1 10

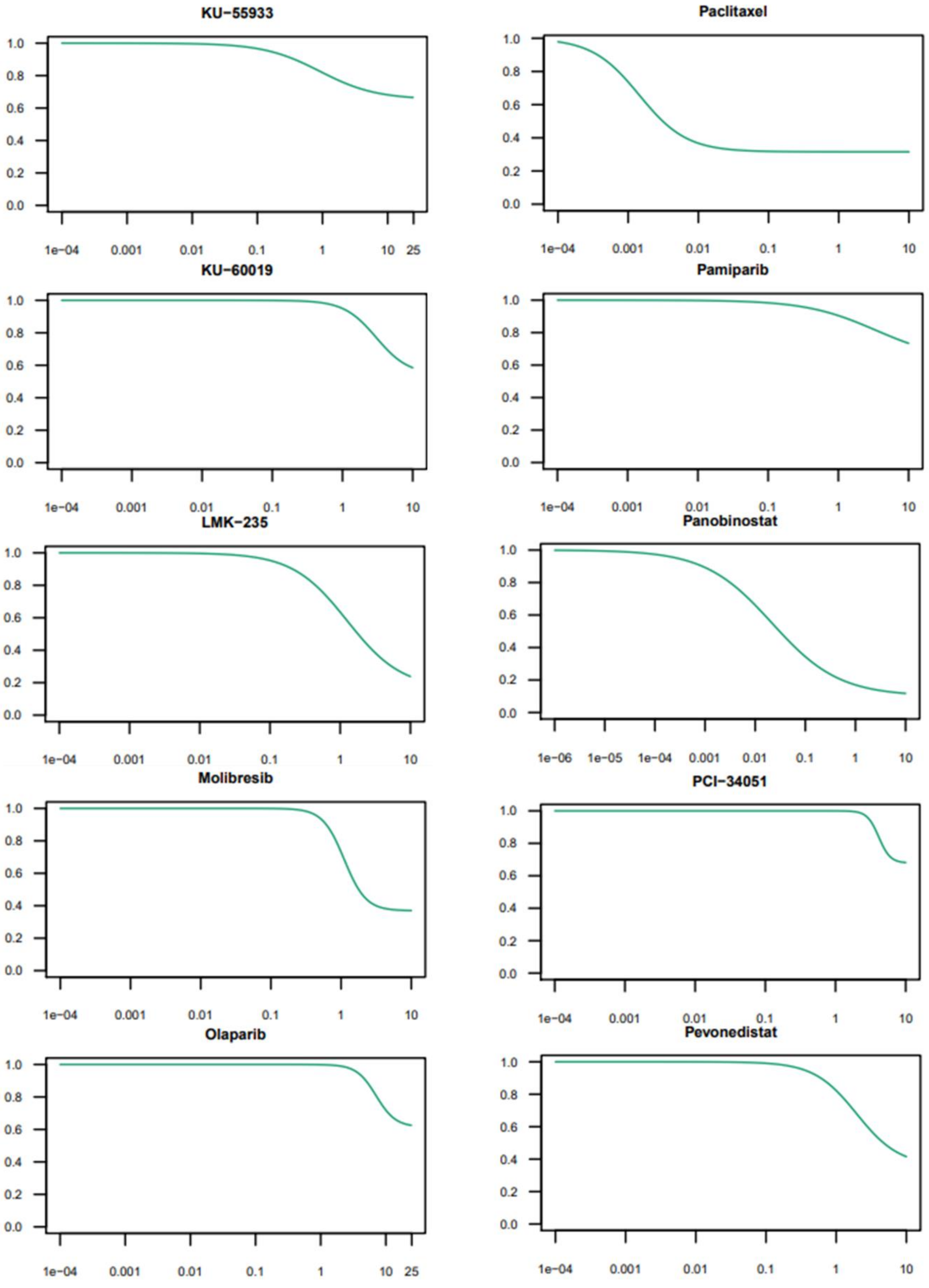
KU−55933
1.0
0.8
0.6
0.4
0.2
0.0
1e−04
0.001
0.01
0.1
1
10
25
Paclitaxel
1.0
0.8
0.6
0.4
0.2
0.0
1e−04
0.001
0.01
0.1
1
10
KU−60019
1e−04
0.001
0.01
0.1
1
10
Pamiparib
1e−04
0.001
0.01
0.1
1
10
LMK−235
1e−04
0.001
0.01
0.1
1
10
Panobinostat
1e−06
1e−05
1e−04
0.001
0.01
0.1
1
10
Molibresib
1e−04
0.001
0.01
0.1
1
10
PCI−34051
1e−04
0.001
0.01
0.1
1
10
Olaparib
1e−04
0.001
0.01
0.1
1
10
25
Pevonedistat
1e−04
0.001
0.01
0.1
1
10

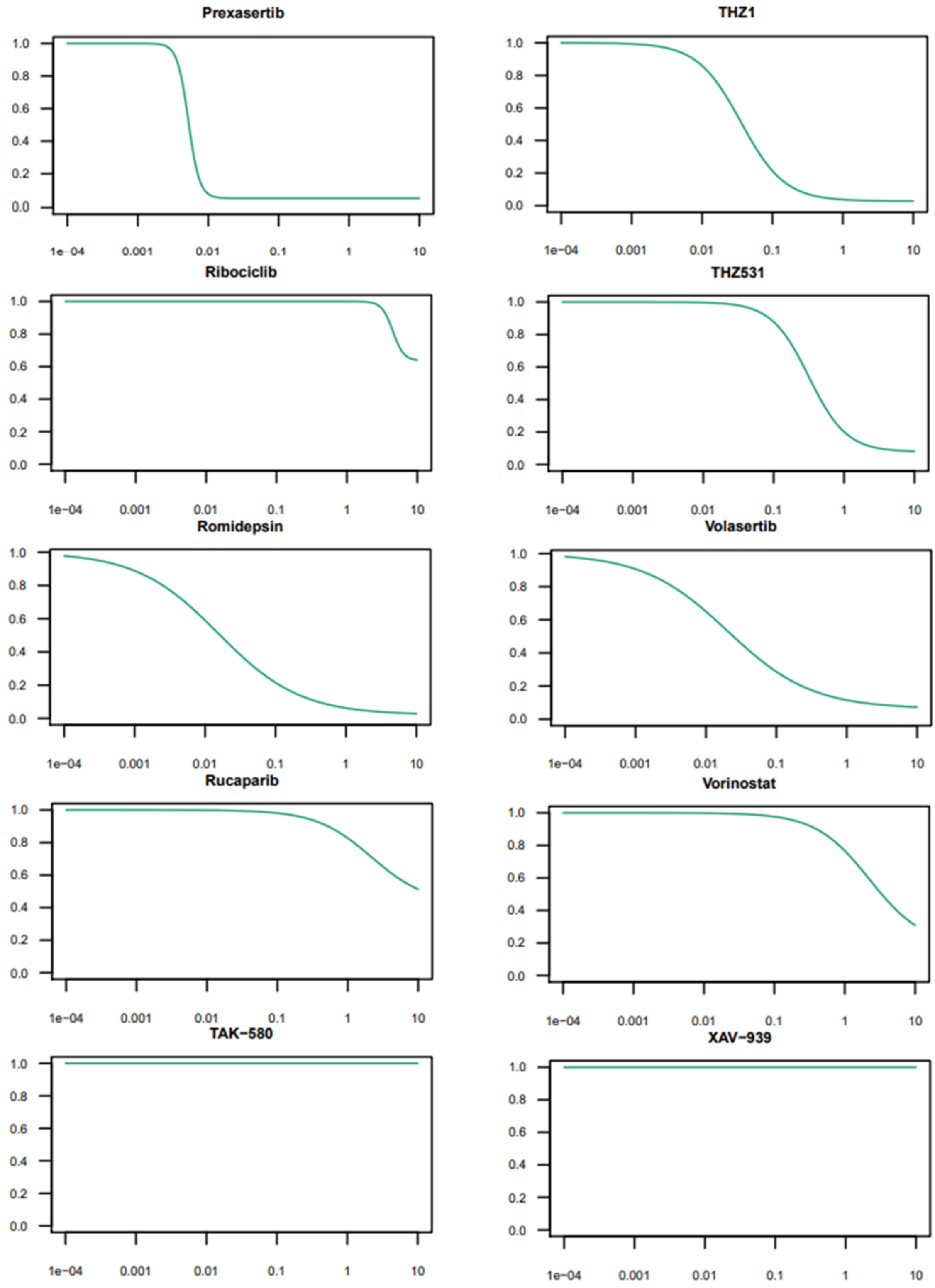


Supplementary figure 2: Individual MTT assay cell viability curves of the compounds used on HEK293T-eGFP cells. IC50 calculations are summarized in Table 1.

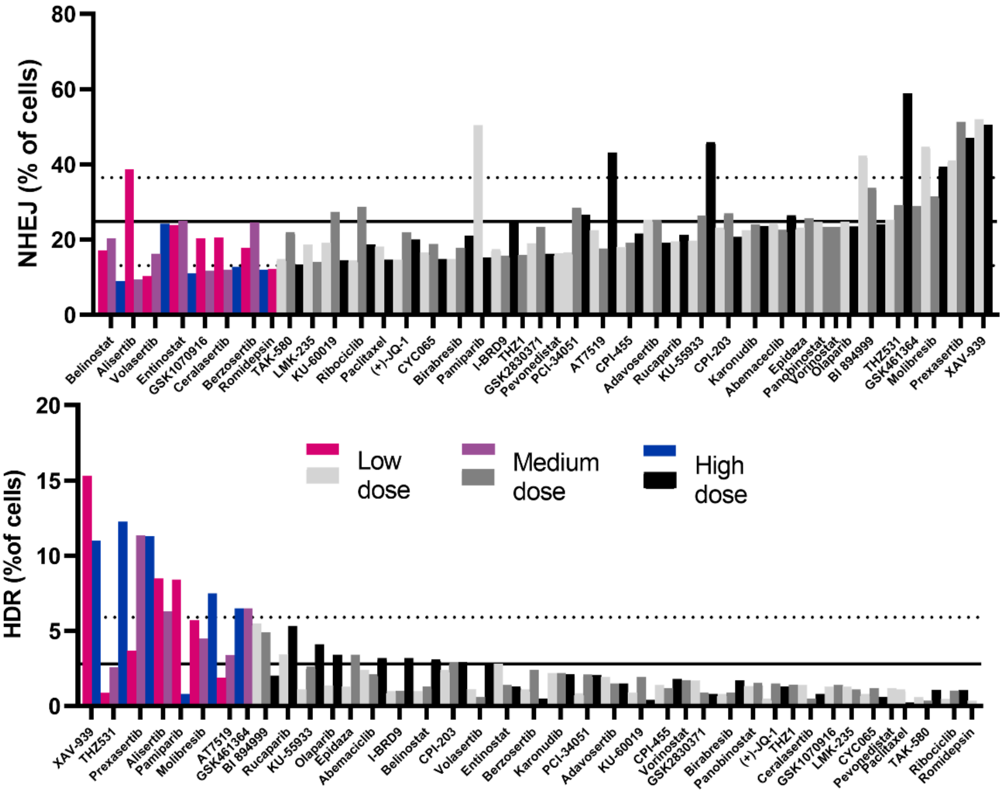


Supplementary figure 3: Effect of the 40 screened compounds on absolute incidence of NHEJ (above, sorted low NHEJ-high NHEJ) and HDR (below, sorted high-low HDR) compared to DMSO-treated controls (mean+- SD as solid and dotted lines, n=29 wells). Colored hits were either lower than mean – SD for NHEJ suppression, or higher than mean + SD for HDR enhancement.

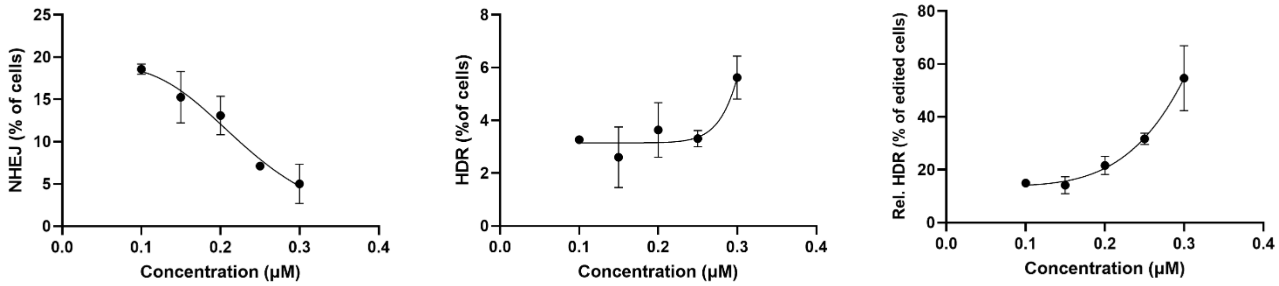


Supplementary figure 4: Effect of a narrow dose range of alisertib on (from left to right) NHEJ, HDR and relative HDR incidences.

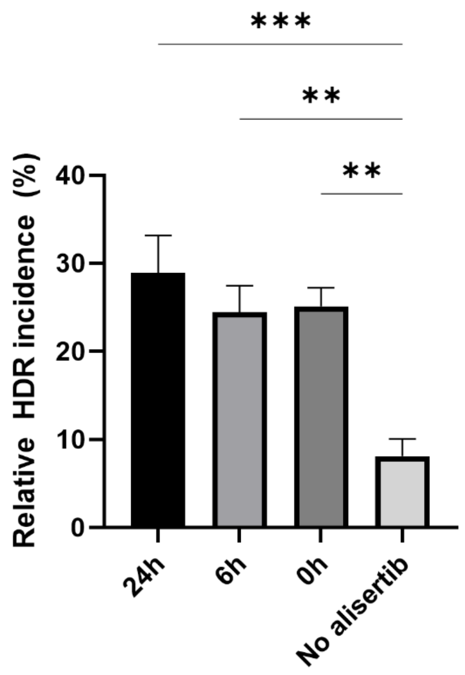


Supplementary figure 5: Alisertib pre-incubation time variation which reveals that simultaneous incubation of alisertib and CRISPR formulations was effective and pre-incubation with alisertib before CRISPR formulations was not significantly more effective than simultaneous incubation.

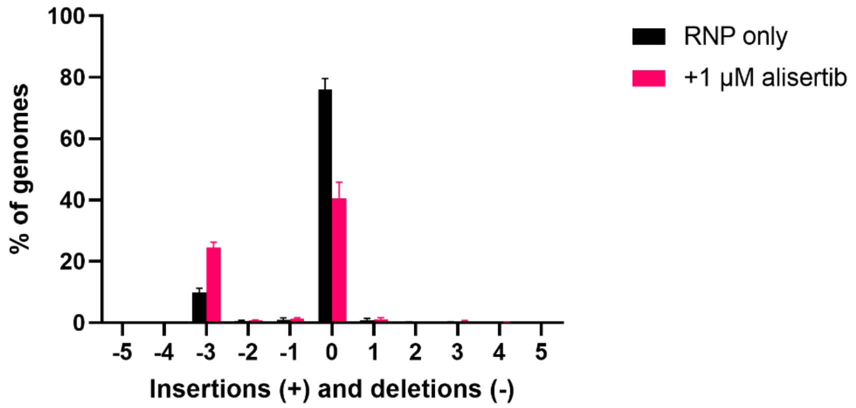


Supplementary figure 6: Mutation distribution found in TIDER analysis, cropped at +- 5 nt. Relatively speaking, most mutations showed up as deletions of a whole codon (-3).
